# CyLaKS: the Cytoskeleton Lattice-based Kinetic Simulator

**DOI:** 10.1101/2021.03.31.437972

**Authors:** Shane A. Fiorenza, Daniel G. Steckhahn, Meredith D. Betterton

## Abstract

Interaction of cytoskeletal filaments, motor proteins, and crosslinkers drives important cellular processes including cell division and cell movement. Cytoskeletal networks also undergo nonequilibrium self-organization in reconstituted systems. An emerging problem in cytoskeletal modeling and simulation is spatiotemporal alteration of the dynamics of filaments, motors, and associated proteins. This can occur due to motor crowding and obstacles along filaments, motor interactions and direction switching, and changes, defects, and heterogeneity in the filament lattice. How such spatiotemporally varying cytoskeletal filaments and motor interactions affect their collective properties is not fully understood. We developed the Cytoskeleton Lattice-based Kinetic Simulator (CyLaKS) for problems with significant spatiotemporal variation of motor or filament properties. The simulation builds on previous work modeling motor mechanochemistry into a simulation with many interacting motors and/or associated proteins. CyLaKS also includes detailed-balance in binding kinetics and movement and lattice heterogeneity. The simulation framework is flexible and extensible for future modeling work. Here we illustrate use of CyLaKS to study long-range motor interactions, filament heterogeneity, motion of a heterodimeric motor, and how changing crosslinker number affects filament separation.

## I. INTRODUCTION

The cellular cytoskeleton performs important biological roles [1], including mitosis [2] and cytokinesis [3] in cell division and cell movement [4, 5]. Key cytoskeletal ingredients include filaments, motor proteins, and other associated proteins. Actin and microtubules are the best-studied filaments [1]. Assemblies of actin and myosin motors participate in cytokinesis [3, 6], cell motility [7–9], and muscle contraction [10–12]. Networks of microtubules and kinesin and dynein motors function in mitotic spindle assembly [2, 13–16] and chromosome segregation [17–20], and beating of cilia and flagella [21–23]. The remarkable ability of the cytoskeleton to dynamically reorganize and exert force is not fully understood [1, 4]. A challenge for theory and simulation is to understand how interactions of filaments, motors and associated proteins can lead to the variety of cytoskeletal structures and dynamics found in cells and reconstituted systems [24–34].

A particular challenge in cytoskeletal theory and modeling is how to study motor and crosslinker behavior that is spatiotemporally altered. For example, kinesin motor activity can be alterered in dense regions along microtubules [35, 36], in ways that differ for different types of motors [36]. These effects may in part be due to short-range interactions between motors [37, 38]. Patchy obstacles created by other proteins binding to the microtubule lattice can alter kinesin and dynein movement [39]. Kinesins that regulate microtubule length and dynamics typically show altered motility at microtubule ends [40–49]. Some kinesin-5 motors can switch their direction of motion along microtubules [50–54], an effect that is not well understood but may be regulated by crowding on the microtubule lattice [54]. Kinesin-1 stepping can be slowed in the presence of crowding molecules in solution, that appear to hinder diffusion of the motor domain [55]. All of these examples illustrate that motor stepping can be altered by the local spatial environment.

In addition, microtubules themselves can have significant spatial and temporal variation. Microtubules can be heterogeneous at the tubulin dimer level if they contain a mixture of tubulin isoforms or are post-translationally modified [56–58]. It appears that motor and non-motor microtubule associated proteins (MAPs) can alter microtubule lattice structure and defects [59–62]. Such changes can alter the binding of kinesin-1 motors [63, 64], an effect that might be explained by elastic anisotropy [65]. Our recent work found that long-range interactions between kinesin-4 motors can explain both changes in motor processivity and velocity at low density and dense accumulation of motors and microtubules ends [34]. How spatiotemporally varying cytoskeletal filaments and motor interactions affect collective motor behavior remains incompletely understood.

Several methods and packages for molecular simulation of filaments and motors are widely used, including Cytosim [66], MEDYAN [67], AFINES [68], and others [69]. Here we build on these methods and develop the Cytoskeleton Lattice-based Kinetic Simulator (CyLaKS)[70]. CyLaKS is designed to facilitate modeling and simulation of cytoskeletal filaments (represented as long, thin rods with a lattice of binding sites for motors and other proteins), motor proteins, and other filament-binding proteins for problems in which spatiotemporal variation of motor or filament properties is significant. CyLaKS uses kinetic Monte Carlo-Brownian dynamics (kMC-BD) methods [71–79]. A primary goal in developing CyLaKS was to build on previous work modeling the mechanochemical cycle of individual motor proteins [80–86] by incorporating the motor ATP hydrolysis cycle in a simulation in which many motors can interact along filaments. Therefore, we extended current simulation packages that incorporate multiple motors to include a more detailed motor model. Explicitly modeling motor mechanochemistry allows results such as nontrivial force-velocity and force-processivity relations [87]. In addition, we implemented detailed-balance in binding kinetics and movement for a physically motivated treatment of force-dependent protein binding/unbinding and diffusion. Finally, we have structured the simulation in a flexible, extensible framework, making it straightforward to elaborate the model to include spatially varying effects such as heterogeneity in the lattice and short- and long-range interactions between motors.

Here we describe the model ingredients and simulation techniques, then illustrate how CyLaKS can be used to study long-range motor interactions, filaments with heterogeneous lattice, motion of a heterodimeric motor, and filament separation due to force from crosslinkers.

## II. MODEL AND SIMULATION

Building on our previous work, CyLaKS implements Brownian dynamics to model physical movement of filaments and kinetic Monte Carlo to model state change (such as protein binding and unbinding) and chemical reactions (such as ATP hydrolysis) [71–79]. Each simulation time step includes first a kinetic Monte Carlo (kMC) substep, then a Brownian dynamics (BD) substep. We implement kinetic Monte Carlo events with a hybrid tau-leaping algorithm [88], which samples from the binomial and Poisson distributions to predict the number of events that will occur in a timestep. The binomial distribution is used for events with a constant probability throughout the simulation, such as ATP hydrolysis in a motor or protein binding (these probabilities are constant in time and identical for every site in the absence of long-range coupling) [72]. For events with a probability that varies, we sample the Poisson distribution. This can occur, for example, when protein unbinding is force dependent, or if long-range motor coupling gives different binding kinetics to each lattice site. We compute the pairwise partition function of each object that an event can act on to calculate the average number of events in the timestep [72]. We then sample the Poisson distrubution to choose the number of events that occur.

In CyLaKS, events are executed in random order on randomly selected members of the population. Multiple events can affect a single population, e.g., binding of the second head and unbinding of the first head for a protein with one head bound. The code enforces that no two events can act on the same object in the same time step, a good approximation if the time step is sufficiently small. During the kMC substep, we implement the following algorithm:

1. Loop over all active objects and sort into appropriate populations.
2. If appropriate, check equilibration status of each protein species. This is done by finding the average number of proteins bound for each species in a time window *t*_c_. If the change in number bound between two windows is less than their standard deviations added in quadrature, the protein species is considered equilibrated. Once all species are equilibrated, the simulation is considered equilibrated.
3. Sample the appropriate statistical distribution for each possible event. If the event has a constant probability, use the binomial distribution. If the event has a probability that can change in space or time, determine the total probability and sample the Poisson distribution.
4. If two or more events target the same molecule, discard at random until only one remains. The probability of each event determines the relative weight when sampling to discard.
5. Execute each event in a random order on random molecules from the appropriate population.

During the Brownian Dynamics substep the filament positions are updated. Due to the relatively large net force that can act on filaments, we run N_BD_ iterations of the BD substep for every kMC iteration that runs to reach filament mechanical equilibrium given the positions of bound motors/crosslinker. To do this, we use a smaller time step *t*_BD_ = *dt/N*_BD_. Our BD substep implements the following algorithim:

1. Update associated protein positions.
2. Sum to find the systematic (non-random) net force and torque acting on each filament.
3. Calculate the translational displacement of each filament’s center of mass due to the systematic force. Movement is projected into parallel and perpendicular directions relative to the long axis of the filament.
4. Calculate the rotational displacement of each filament’s orientation vector due to the systematic torque. The direction of this displacement is always perpendicular to the long axis of the filament.
5. Calculate the random displacements of center of mass and random reorientation due to thermal noise.
6. Apply the displacements and update filament positions.

The simulation continues to alternate kMC and BD substeps until it has reached the specified total run time.

CyLaKS is written in C/C++ and uses a combination of templated functions and class inheritance for modularity while retaining reasonable performance. Basic molecular ingredients such as binding and catalytic heads, linear and angular springs, and discrete binding sites all inherit from a common Object class. This allows us to take advantage of class polymorphism, a property of C++ classes that allows containers such as vectors or arrays to hold different data structures, as long as they inherent from each other. Additionally, we can define the same function differently for different inheritance classes. By using these two properties together, we can hold all active molecular species in the same container and iterate while calling a generic update function that act appropriately on the species it’s called for. This structure decouples the main kMC-BD algorithim from the details of the species. This allows addition of new species or mechanisms without modifying the majority of the code. Furthermore, classes that are specialized versions of others inherit directly from the more general class; for example, a catalytic head is a specialized version of a binding head. This design allows us to re-use common code between classes while adding additional ingredients needed by the specialized class. As a result, code only needs to be written once even if re-used, and code readability and validation are improved.

To increase overall performance, proteins in CyLaKS are not explicitly modeled while in solution. They are instead drawn from a reservoir list when they first bind to an object. Additionally, unbound springs and/or unbound heads of proteins with only one of their heads bound to a filament, are not explicitly modeled. Finally, all interactions (springs, long-range potentials, etc) have a cutoff distance, to avoid modeling low-probability events.

### A. Filaments

The current filament model in CyLaKS is based on a microtubule idealized as a rigid single protofilaments where each tubulin dimer corresponds to a discrete binding site on a 1-D lattice (Fig. 1a). In the future, extending the model to multiple protofilaments or actin filaments would be straightforward. We implement the Brownian dynamics algorithm of Tao et al. [89], as in our previous work [71–79]. The microtubule center-of-mass position 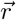 evolves according to

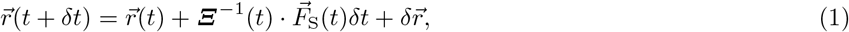

where ***Ξ***^−1^ is the inverse friction tensor of the filament, 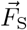 is the deterministic (systematic) force acting on the filament, *k_B_T* is the thermal energy, and *δt* is the time step. Diffusion occurs due to the random displacement 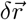 which is anisotropic and Gaussian-distributed with variance 2*k_B_T* ***Ξ***^−1^(*t*)*δt*. The inverse friction tensor ***Ξ***^−1^(*t*) is

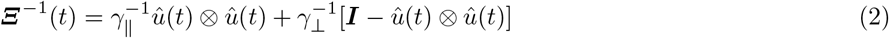

where *γ*_‖_ and *γ*_⊥_ are the translational friction coefficients for motion parallel and perpendicular to the axis of the filament, respectively, and 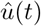 is the filament orientation at time *t*. The operators 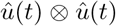 and 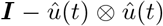 project onto directions that are parallel and perpendicular to the filament, respectively.

**FIG. 1.**
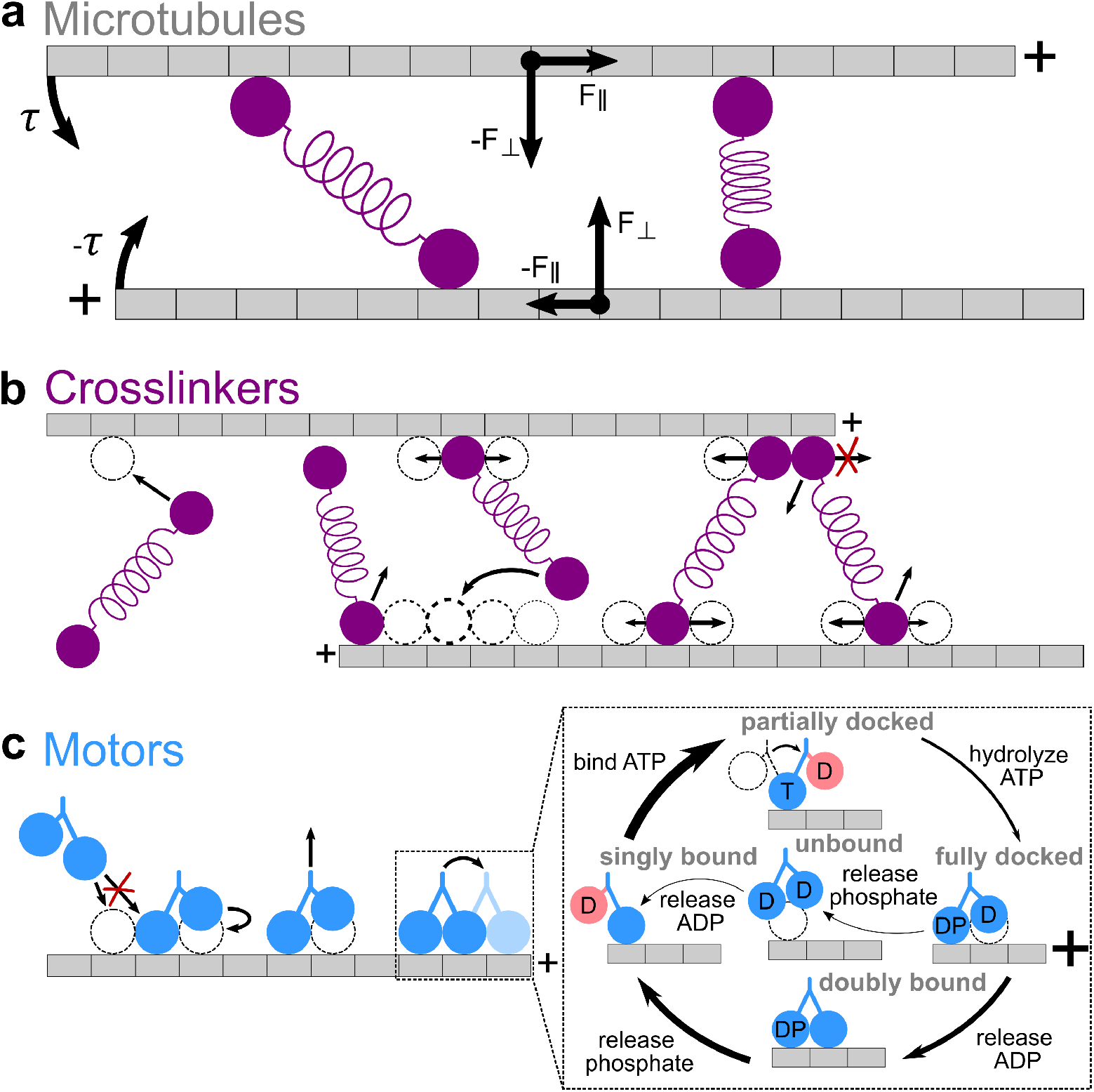
CyLaKS model ingredients A. Microtubules. Microtubules are modeled as single protofilaments, where each tubulin dimer corresponds to a discrete site on a 1-D lattice. Each microtubule has a plus- and minus-end. Associated proteins exert force and torque on filaments, causing 2-D translaton and rotation about each filament’s center of mass. B. Crosslinkers. Each crosslinker head can independently bind to, unbind from, and diffuse on the filament lattice. The relative probability of the second head binding to each sites is represented by dotted circles of different thickness. The relative probability of heads diffusing is represented by arrow length. Steric exclusion prevents more than one crosslinker head from occupying the same binding site. Crosslinker heads cannot diffuse off filament ends. C. Motor proteins. Motors can bind to, unbind from, and step toward the plus-ends of filaments. Inset, mechanochemical cycle. Motor heads can be bound to ADP (D), ATP (T), ADP·Pi (DP), or nothing (empty). Red coloring labels head which cannot bind to the microtubule due to necklinker tension. Arrow thickness represents the relative probability of each transition. Steric exclusion prevents more than one motor head from occupying the same binding site.

Filament rotation is described by the time evolution of the orientation vector 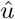,

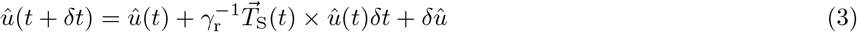

where *γ*_r_ is the orientational friction coefficient of the filament, 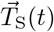 is the deterministic torque on the filament about its center of mass at time *t*, and 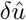 is a random reorientation that is anisotropic and Gaussian-distributed with variance 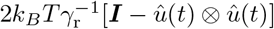.

The filament friction coefficients are

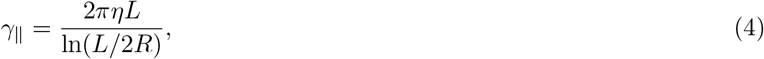

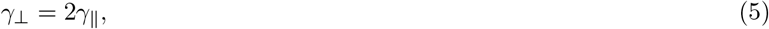

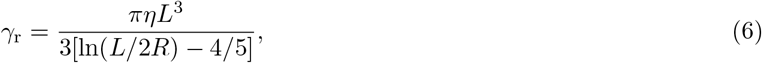

where *η* is the viscosity of the fluid, *L* is the length of the filament, and *R* is the radius of the filament [90].

### B. Binding kinetics

Motor and crosslinker binding to filaments has an associated free-energy change, which includes the binding energy and the energy in a crosslinker spring if it is stretched or compressed. The energy change upon binding we denote Δ*U*_0_ = *U*_bound_ − *U*_unbound_. Then the association (binding) and dissociation (unbinding) rates can be written

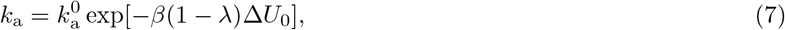

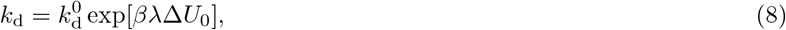

where *β* = 1/*k*_B_*T* is the inverse thermal energy and *λ* a dimensionless constant typically with 0 ≤ *λ* ≤ 1. Here *λ* weighs how strongly the interaction affects binding or unbinding. We typically assume *λ =*1/2 in the absence of experimental measurement of this value.

The change in free energy upon binding can depend on the state of the system: for example when proteins co-operatively bind, the effective free energy of the bound state changes if nearby proteins are also bound. If a state change such makes the new binding-energy difference *U*_bound_ − *U*_unbound_ = Δ*U*, the association and dissociation rates change accordingly. As an example, consider an attractive cooperative interaction that lowers the free energy of the bound state. This will lower Δ*U* relative to Δ*U*_0_ in Eqs. 7 and 8, resulting in the association rate *k*_a_ increasing and dissociation rate *k*_d_ decreasing.

### C. Crosslinkers

Non-motor crosslinking proteins are modeled as two independent binding heads connected by a spring (Fig. 1b). We assume that each head binds to filaments separately, so crosslinkers bind one head (entering the one-head-bound or singly bound state) before the second (entering the two-heads-bound or crosslinking state). As currently implemented, we require that the second head cannot bind to the same filament as the first.

The first heads bind to filaments at rate *Nk*_on_*c*_bulk_, where *N* is the number of unoccupied sites on the filament, *k*_on_ is the per-site crosslinker association rate, and *c*_bulk_ is the concentration of unbound crosslinkers in solution. When one head is bound, crosslinker heads can unbind at rate *k*_off,1_, or diffuse to adjacent unoccupied sites at rate 2*D*_1_*/*Δ^2^, where *D*_1_ is the one-head-bound diffusion coefficient and Δ the length of a lattice site (approximately 8 nm for tubulin dimers).

One-head-bound crosslinkers can form crosslinks to adjacent microtubules by binding the second head at a bare rate *k*_on_*c*_eff_, where *c*_eff_ is the effective concentration of the second head. In most cases this effective concentration is not experimentally measured. Therefore, we use a model in which the second head can sample the volume of a half-sphere with radius *r*_0_ centered around the first head when singly bound, where *r*_0_ is the length of the crosslinker. Once a crosslink forms, the crosslinker spring energy is

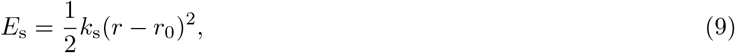

where *k*_s_ is the spring constant, *r* the length of the crosslinker as currently bound, and *r*_0_ the unperturbed length. We take this change in energy into account as discussed above (Eqs. 7 and 8), giving

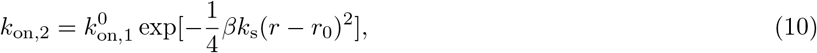

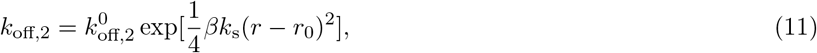

where we have assumed *λ* = 1/2. We typically assume that 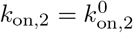, i.e., the biochemistry of binding heads does not change when one or two heads are bound. It is possible to model mechanisms that may alter the off rate when two heads are bound. Crosslinker heads unbind from the microtubule independently, with rate *k*_off,2_ given by Eq. 11.

Crosslinker heads diffuse independently with a diffusion coefficient *D*_2_ that is scaled by the appropriate Boltzmann factor for the change in spring extension during a diffusive step. A diffusive step that stretches/compresses the spring is energetically unfavorable and occurs at lower rate, while a diffusive step that relaxes the spring stretch/compression is energetically favorable and occurs at higher rate,

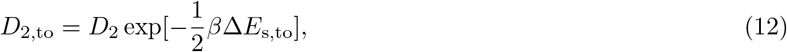

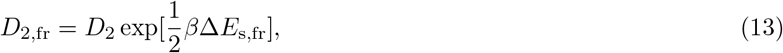

where Δ*E*_s,to_ denotes a change in energy of a step that relaxes the spring, and Δ*E*_s,fr_ is the change in energy of a step that extends/compresses the spring.

Steric interactions prevent two crosslinker heads from occupying the same lattice site.

### D. Motors

We choose the motor model in CyLaKS to allow studies of many interacting motors while also including a motor mechanochemical cycle [80–86]. We implemented motors as two coupled catalytic heads. Each head can bind to and unbind from the filament as a passive binding head would. However, they can also bind a ligand and transition through a nucleotide hydrolysis cycle based on the kinesin-1 stepping cycle (Fig. 1c) [87, 91–96]. Details of this cycle may be differ between motor species, but any basic mechanochemical cycle that facilitates asynchronous binding and unbinding of two binding heads can lead to similar stepping [93, 97].

In the currently implemented motor model, motors bind similarly to crosslinkers at rate *Nk*_on_*c*_bulk_, where *N* is the number of unoccupied sites on the microtubule, *k*_on_ the association rate, and *c*_bulk_ the concentration. For simplicity, we assume that the first motor head to bind is always the leading head (for plus-end-directed motors, closer to the plus-end). The biochemical states possible for each motor head are bound to ATP, APP, ADP·Pi, or no ligand (empty).

We assume that any motor not attached to a microtubule has both motor heads bound to ADP. Upon binding to the microtubule, the motor head releases ADP, becoming empty. While empty or ATP-bound, motor heads are strongly bound and cannot unbind from the microtubule. ATP binding to the empty head at rate *k*_ATP_ induces a conformational change that swings the second (unbound) head forward, partially docking the motor. The first (bound) head then hydrolyzes ATP to ADP·Pi at rate *k*_hydro_, which fully docks the motor. Next, the motor can either unbind its first (bound) head at rate *k*_off,1_, which terminates its run, or bind its second (unbound) head at rate *k*_on_*c*_eff_, continuing its run. The ratio between *k*_off,1_ and *k*_on_*c*_eff_ determines how many steps occur on average before the motor unbinds. While the motor is doubly bound, the rear head unbinds with rate *k*_off,2_ and the front head cannot unbind. Upon rear head unbinding, the motor has completed one ATP hydrolysis cycle and moved forward one site.

Both the run length and velocity of motors depend on the force being applied, e.g., from an optical trap or molecular load [87]. Andreasson et al. found that this force dependence could be properly accounted for by a 3-state model, where ATP binding and hydrolysis is condensed into one step. The rate of ATP binding while in the singly bound state is modified as

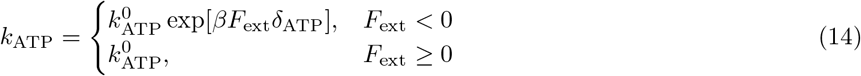

where *F*_ext_ is the external force acting on the motor and *δ*_ATP_ is a distance parameter that controls the strength of force dependence. External force is negative if applied in the opposite direction of motor stepping and positive if in the stepping direction. The rate of unbinding while in the fully docked state is modified as

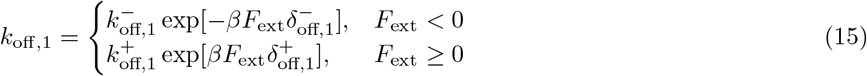

where ± superscripts correspond to assisting and hindering loads, respectively. In this form, forces always act to enhance the unbinding rate of motors while in the docked state. However, this enhancement does not necessarily have to be symmetric. In our simulations, we typically set 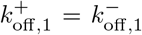 and 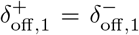. Finally, we also modify the rear-head unbinding rate of doubly bound motors as

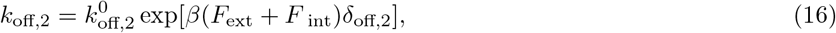

where *F*_int_ is the internal necklinker tension of the motor, estimated to be ≈ 26 pN for kinesin-1 [87].

### E. Cooperative binding and movement

CyLaKS can model both short- and long-range cooperative interactions between motors and crosslinkers bound to a filament [34]. Based on previous work [38], we implemented a model of short-range binding cooperativity as an attractive nearest-neighbor interaction. The interaction energy when one motor (or crosslinker) head binds near an adjacent bound head is −*ϵk*_B_*T*, assuming an interaction range of one lattice site. When implementing this interactions, we set *λ* = 1, leading to

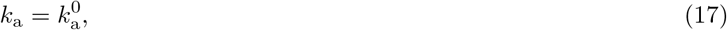

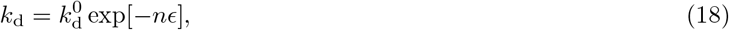

where *n* = 0, 1, 2 is the number of neighbors. As currently implemented in CyLaKS, each type of protein (crosslinkers and motors) can have a different value of *ϵ* for interactions with proteins of the same type or different types. We assume that binding heads belonging to the same protein (for example, the two binding heads of a motor) do not interact with each other.

As currently implemented, we model long-range binding cooperativity between proteins with an attractive potential *E*(*x* − *x_m_*), where

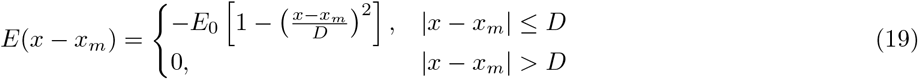

which is negative up to the cutoff distance *D* and zero beyond that distance. Here *x* is the distance along lattice, *x_m_* is the position of the motor that is inducing the interaction, *E*_0_ is the strength of the interaction energy, and the curvature of the potential is determined by *E*(*D*) = 0. Currently we assume that the interaction induced by multiple motors can superpose up a maximum *E*\*. In modeling long-range cooperativity we typically set *λ* = 1/2, giving

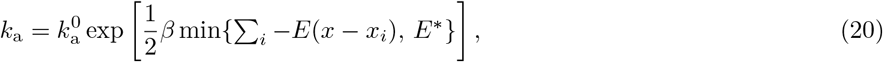

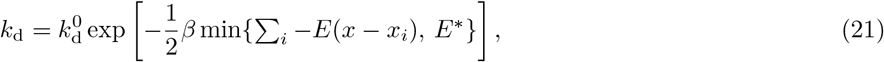

where the sum is over bound motors.

The long-range attractive interaction (Eq. 19) can affect the motor stepping cycle. As currently implemented, we assume that the internal necklinker tension that couples motor heads is decreased by the interaction energy. This is one possible mechanism by which an interaction could affect motor stepping. However, other mechanisms are also possible. We extended the model of necklinker tension during motor stepping of Andreasson et al. [87] by introducing a new doubly bound to partially docked pathway with rate

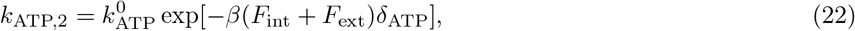

where *δ*_ATP_ is a distance parameter that controls how strongly front-head ATP binding is affected by force. If ATP binds to the front head, the rear head detaches from the microtubule and swings forward, skipping the singly bound state. Using parameters previously estimated for kinesin-1 (Table I), the doubly bound off- and ATP-binding rates are 2375 and 1.1810 × 10^−9^ s^−1^, respectively. If the necklinker tension is reduced by the long-range interaction (Eq. 19), the doubly bound ATP binding rate can become significant. We implement this reduction of tension by multiplying the doubly bound ATP binding and motor off rates by exp[*β*/2 min{Σ*_i_* −*E_i_*, *E*\*}] and exp[− *β*/2 min{Σ_*i*_ −*E_i_*, *E*\*}], respectively.

**TABLE I.**
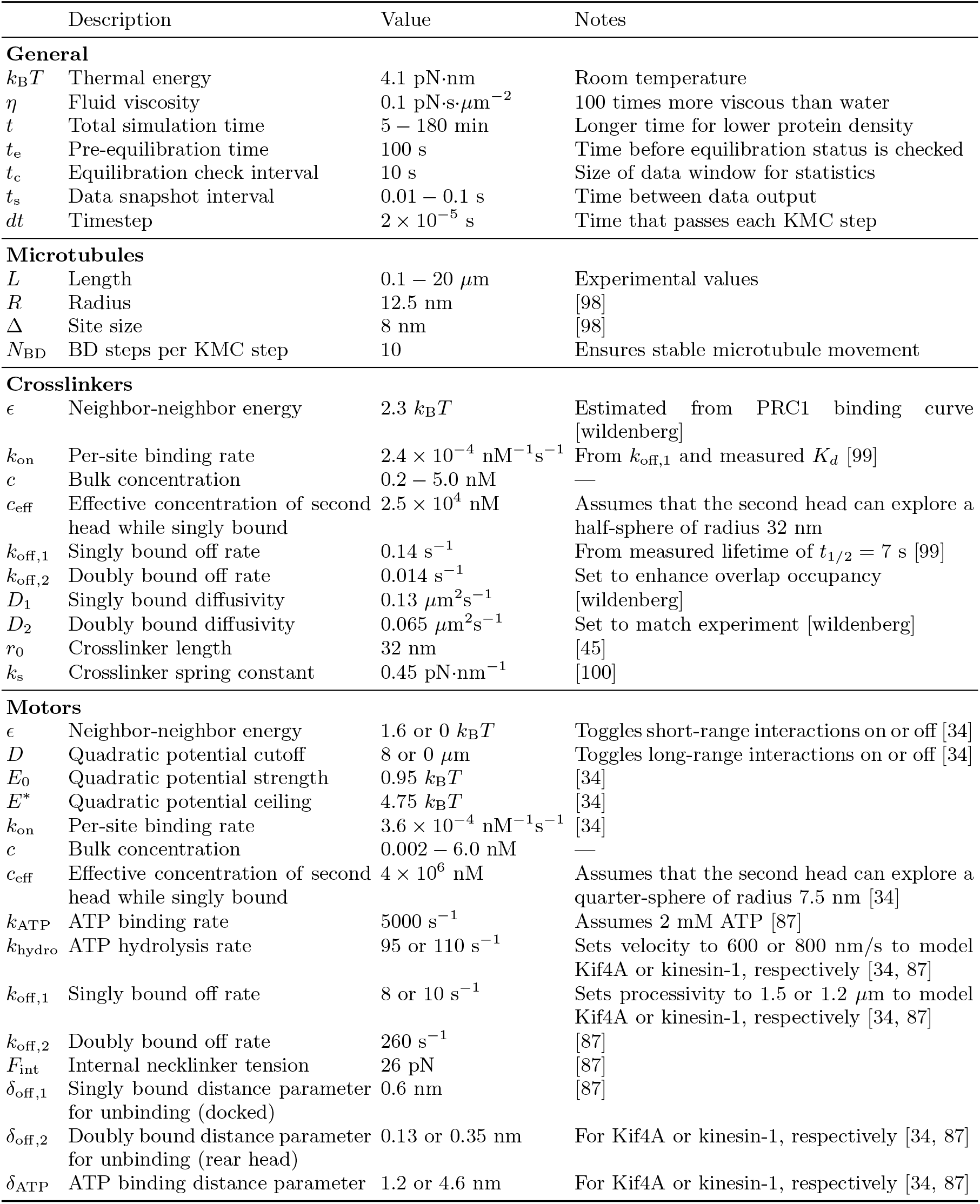
Parameters used in CyLaKS.

### F. Parameters

Parameters and their source are summarized in table I. Here we discuss the estimation of parameters not directly constrained by experiments. We estimate *c*_eff_ for crosslinkers by calculating the volume of a half-sphere of radius *r*_0_. The effective concentration is that of one molecule in this volume. With *r*_0_ = 32 nm,

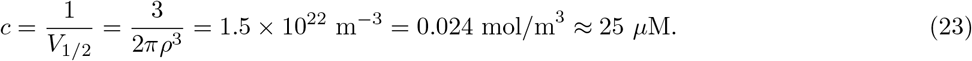

For motors, we assume each motor head can only explore a quarter-sphere, and we use *r*_0_ = 7.5 nm as the linkage length. This gives *c*_eff_ ≈ 4 mM.

The off-rate of crosslinking PRC1 heads has not been experimentally measured. However, it has been observed that PRC1 binds at least 28 times more strongly to anti-parallel microtubule overlaps versus single or parallel microtubules [101]. The effective concentration increase due to binding (summarized in *c*_eff_) partially accounts for this. However, we found that decreasing *k*_off,2_ by a factor of 10 compared to *k*_off,1_ could give the overlap enhancement found experimentally.

Both the one-head-bound and crosslinking diffusion coefficient of PRC1 have been measured experimentally [102]. We directly use the value for singly bound heads. To determine the diffusion coefficient when crosslinking, we take into account the effects of force dependence into account. A simulation diffusion coefficient of 0.065 *μ*m^2^s^−1^ leads to an effective crosslinking diffusion coefficient of 0.021 *μ*m^2^s^−1^, as measured [102].

For motors, the hydrolysis rate *k*_hydro_ is determined by the average velocity expected for the motor because ATP hydrolysis is the rate-limiting step of the mechanochemical cycle (Fig. 1c). For Kif4A, a hydrolysis rate of 95 s^−1^ yields an average velocity of 600 nm/s. For kinesin-1, a hydrolysis rate of 110 s^−1^ yields an average velocity of 800 nm/s. The value of *k*_off,1_ is controls the processivity because motors can only unbind while in the fully docked state. For model Kif4A, a singly bound off rate of 8 s^−1^ yields an average processivity of 1.5 *μ*m. For kinesin-1, a singly bound off rate of 10 s^−1^ yields an average processivity of 1.2 *μ*m.

For the Kif4A model, we fit the parameters of the short- and long-range interaction to data on low-density Kif4A motility. We implemented this using the nonlinear least-squares optimization routine from the SciPy python library [103]. The fit estimated the six parameters *ϵ*, *D*, *E*_0_, *E*^∗^, *δ*_off,2_, and *δ*_ATP_; final parameters were determined by hand adjustment.

### G. Validation

To validate the simulations, we compared simulation results to theory for microtubule, crosslinker, and motor movement (Fig. 2). For microtubules (Fig. 2a–c), we predicted and measured the diffusion parallel and perpendicular to the filament long axis. In the absence of applied force, the center of mass diffuses with ⟨*x*^2^⟩ = 2*Dt* (Fig. 2b). Under a constant applied external force 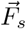, the filament velocity is 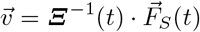 (Fig. 2c).

**FIG. 2.**
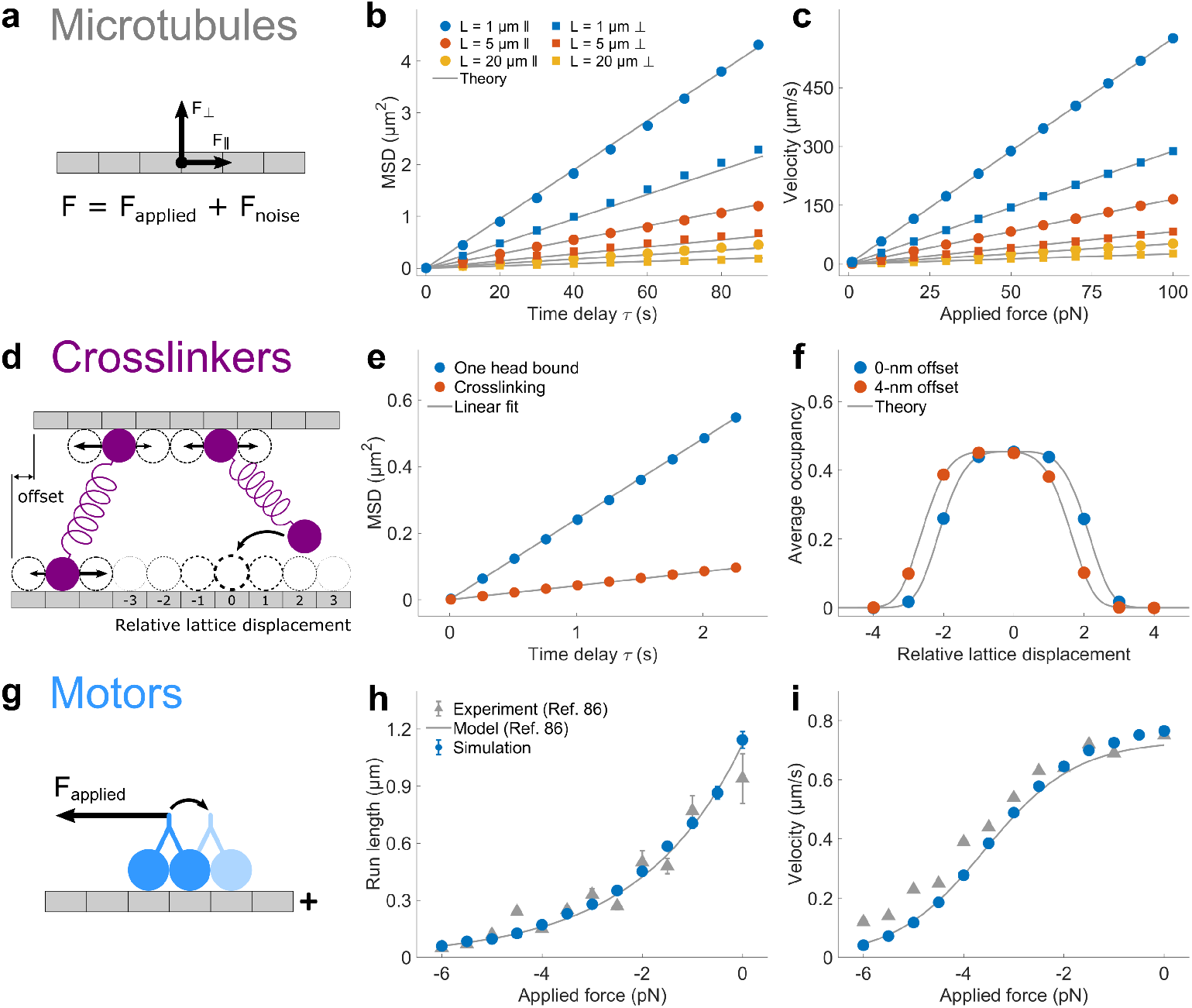
Simulation validation. (A-C), Microtubules. A. Schematic of 2D microtubule movement. B. Plot of mean-squared-displacement (MSD) of microtubule center of mass as a function of time delay *τ* for varying microtubule length, for movement parallel (circles) and perpendicular (squares) to the filament long axis. Theory is the prediction from 1D diffusion. Data were averaged from six independent simulations; error bars show standard error of the mean and are smaller than the points. C. Plot of velocity of microtubule center of mass as a function of applied force for varying microtubule length, for movement parallel (circles) and perpendicular (squares) to the filament long axis. Theory is the prediction from constant-force motion. Data were averaged from six independent simulations; error bars correspond to standard error of the mean and smaller than the points. (D-F), Crosslinkers. D. Schematic. Two microtubules are fixed with a vertical separation of 32 nm, the length of the crosslinkers. The top microtubule is horizontally displaced by the offset distance, where an offset of 0 nm means the lattices of each microtubule are aligned. E. Plot of crosslinker mean-squared-displacement (MSD) versus time delay *τ*. The diffusion coefficient is determined from a linear fit (0.121 *μ*m^2^s^−1^ for one head bound; 0.0213 *μ*m^2^s^−1^ for crosslinking). F. Plot of average second head occupancy versus relative lattice displacement for two different values of the microtubule offset. Theory is the prediction from statistical mechanics (see text). (G-I) Motors. G. Schematic. Motors move under a constant hindering force. Plots of run length (H) and velocity (I) versus applied force. Experimental data and model from previous work [87]. These runs used the kinesin-1 parameter set of Table I.

For crosslinkers, we measured both crosslinker diffusion and formation of a crosslink when one bound head is fixed (Fig. 2d–f). The PRC1 diffusion coefficient when one head is bound matches experimental results [102]. The measured
diffusion coefficient of crosslinking PRC1 is 0.024 ± 0.003 *μ*m^2^s^−1^ [102]. A slightly larger bare simulation diffusion coefficient of 0.0655 *μ*m^2^s^−1^ gives an effective PRC1 diffusion coefficient of 0.021 *μ*m^2^s^−1^ because of force-dependent diffusion (Fig. 2e). For crosslink formation when one head is bound and fixed in place, the occupancy of a site on the other filament is

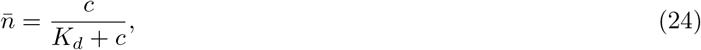

where 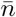 is the average fractional occupancy of the site, *c* is the concentration of the binding head, and *K_d_ = k_d_/k_a_* is the dissociation constant. We use *c = c*_eff_ when measuring the occupancy of the crosslinking head. The dissociation constant varies for each lattice site because of the crosslinker spring extension according to Eqs. 7 and 8, giving

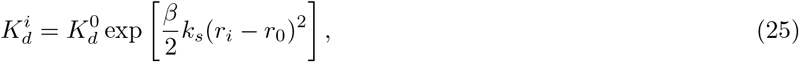

where *r_i_* is the crosslinker extension for binding at site *i*. This matches the simulated occupancy for fixed microtubule position (Fig. 2f).

For motors, we compared to previous measurements and model of kinesin-1 movement under hindering load [87] (Fig. 2g–i). The force dependence of both the run length (Fig. 2h) and velocity (Fig. 2i) in our simulations match previous work. These simulations use the kinesin-1 parameter set (Table I), with a motor concentration of 50 pM and microtubule length of 80 *μ*m, with results averaged over 4 independent simulations.

## III. RESULTS

Here we illustrate the types of simulations that can be done in CyLaKS using four examples. First, we build on our previous work to show how long-range interactions between Kif4A motors can lead to microtubule-length-dependent accumulation at plus-ends [34]. Second, inspired by previous work on heterogeneity in the microtubule lattice [56–58], we show how motor motility changes if the lattice contains a random mixture of sites with different motor binding affinity. Third, building on previous work on heterodimeric kinesin motors [104–109], we create a toy model of a heterodimeric motor in which one head is immobile while bound while the other head can diffuse while singly bound. The model demonstrates the expected crossover from directed to diffusive motility as the diffusion coefficient of the second head increases. Fourth, we demonstrate how varying the number of crosslinkers between two microtubules can change the equilibrium microtubule separation.

### A. Length-dependent end-tags

In recent work, we used CyLaKS to study how collective motor behavior can change due to long-range interactions [34]. Kif4A is a human kinesin-4 motor that accumulates densely on microtubule plus-ends, forming end-tags. Previous work found that end-tag length increases linearly with the length of the microtubule on which they form [45]. This length dependence persists for microtubules up to 14 *μ*m long, even though single Kif4A motors have a run length of 1 *μ*m. We found that this surprising result could not be explained by a conventional motor model with only short-range interactions between motors. A combination of short- and long-range cooperative interactions between motors allowed length-dependent end-tags to form in our model [34] (Fig. 3).

**FIG. 3.**
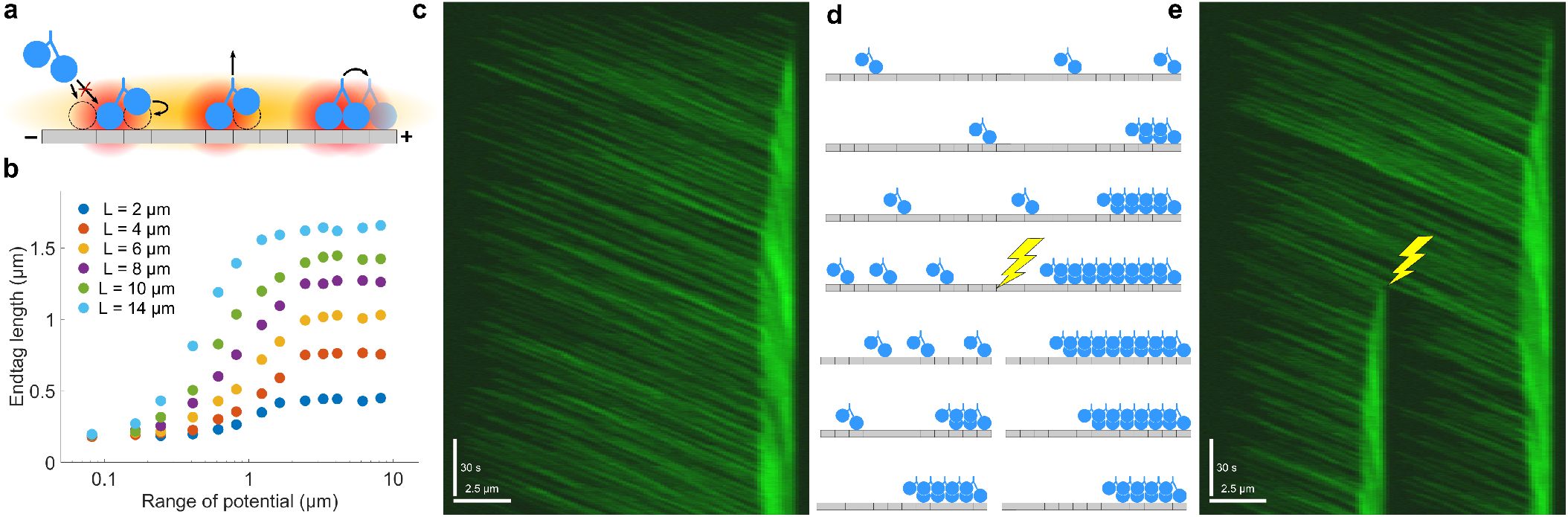
Kif4A end-tag formation due to long-range motor interactions A. Schematic of the motor interaction model. Motors interact by short- and long-range cooperativity, and the long-range interaction affects both binding and stepping. B. Plot of end-tag length versus potential range for varying microtubule length. C. Simulated kymograph of end-tag formation. 67% of simulated motors are fluorescently labeled. Scale bars are 2.5 *μ*m and 30 seconds. D. Schematic of the microtubule ablation simulation. After an end-tag forms, the microtubule is split in half. A new end-tag forms on the new microtubule, while the old end-tag shrinks. E. Simulated kymograph of microtubule ablation simulation. 67% of simulated motors are fluorescently labeled. Scale bars are 2.5 *μ*m and 30 seconds.

Previous work found that bound kinesin-1 motors can increase the binding rate of other motors up to 6 *μ*m away [63]. We therefore implemented a long-range interaction as a Gaussian increase in binding affinity up to a distance *D* (Eqs. 20 and 21). We found that long-range interactions that only affect binding kinetics only partially explain the data. Our experimental results showed that motor velocity significantly decreases as motors bind [34]. To predict this, the long-range interaction in our model must affect motor stepping in addition to binding. Since we lack direct experimental evidence on how the interaction might alter motor mechanochemistry, we modeled one plausible mechanism by which this might happen. We assumed that the long-range attractive potential could reduce the internal tension of the necklinker. This reduces rear-head unbinding from the microtubule and increases front-head ATP binding while doubly bound (Eqs. 16 and 22). We fit *ϵ*, *D*, *E*_0_, and *E*\* to experimental data on Kif4A at low density and found good agreement [34]. Without any changes to these model parameters, our model quantitatively matches the length-dependent end-tags found experimentally at higher motor concentration.

Here we extend our previous work by examining how the range of the motor interaction affects end-tag length (Fig. 3a–e). The interaction range must be at least 1 *μ*m to begin to approach the experimental values (Fig. 3b). We also find that for end-tag formation specifically, length saturates at a range of 2.5 *μ*m. To illustrate the dynamics of end-tag formation, we generated a simulated motor kymograph (Fig. 3c). Our model predicts that breaking a microtubule in two would cause the original end-tag to shrink while a new end-tag forms at the plus-end of the new microtubule (Fig. 3d). Both of these predictions are matched by our simulations (Fig. 3e).

### B. Tubulin heterogeneity

The molecular details of motor and MAP interaction with tubulin affect binding movement, causing different tubulin isomers and post-translational modification to influence protein binding and motor movement [110–112]. Previous work found that differences in the tubulin carboxy-terminal tail (CTT) were associated with a decrease in the run length of kinesin-1 while leaving velocity unaffected [110]. However, these same CTT mutants reduced both the velocity and run length of kinesin-2. These raises the possibility that heterogeneous microtubules made up of a mix of different tubulin isoforms or modifications could cause dimer-specific variation in motor or MAP behavior. To model these effects in CyLaKS, we allow each lattice site to have different interactions with a motor or MAP. Here, we illustrate the effects of specifying site-specific binding affinity *ζ_i_* for site *i*, which scales the dissociation constant for binding as

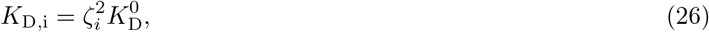

where 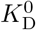 is the original value of the dissociation constant. The parameter *ζ_i_* is squared in this expression because we currently implement it by dividing the binding rate and multiplying the unbinding rate by the same factor.

We then examined the effects of introducing a variable, randomly located fraction of weak-binding sites with *ζ* = 3 for a high-processivity motor (Fig. 4). We varied the fraction of weak-binding sites from 0 to 1. As expected, the average motor run length and lifetime drop as the fraction of weak-binding sites increases, while the motor velocity is unchanged. Remarkably, when just 5% of sites on a 1000-site lattice are weakly binding, the processivity and lifetime of motors are both significantly affected (Fig. 4b). We also see that when all sites are weakly binding, the motor processivity and lifetime decreases by a factor of 9, as expected. Simulated kymographs show a dramatic reduction in motor activity as the fraction of sites with weak binding is increased (Fig. 4c).

**FIG. 4.**
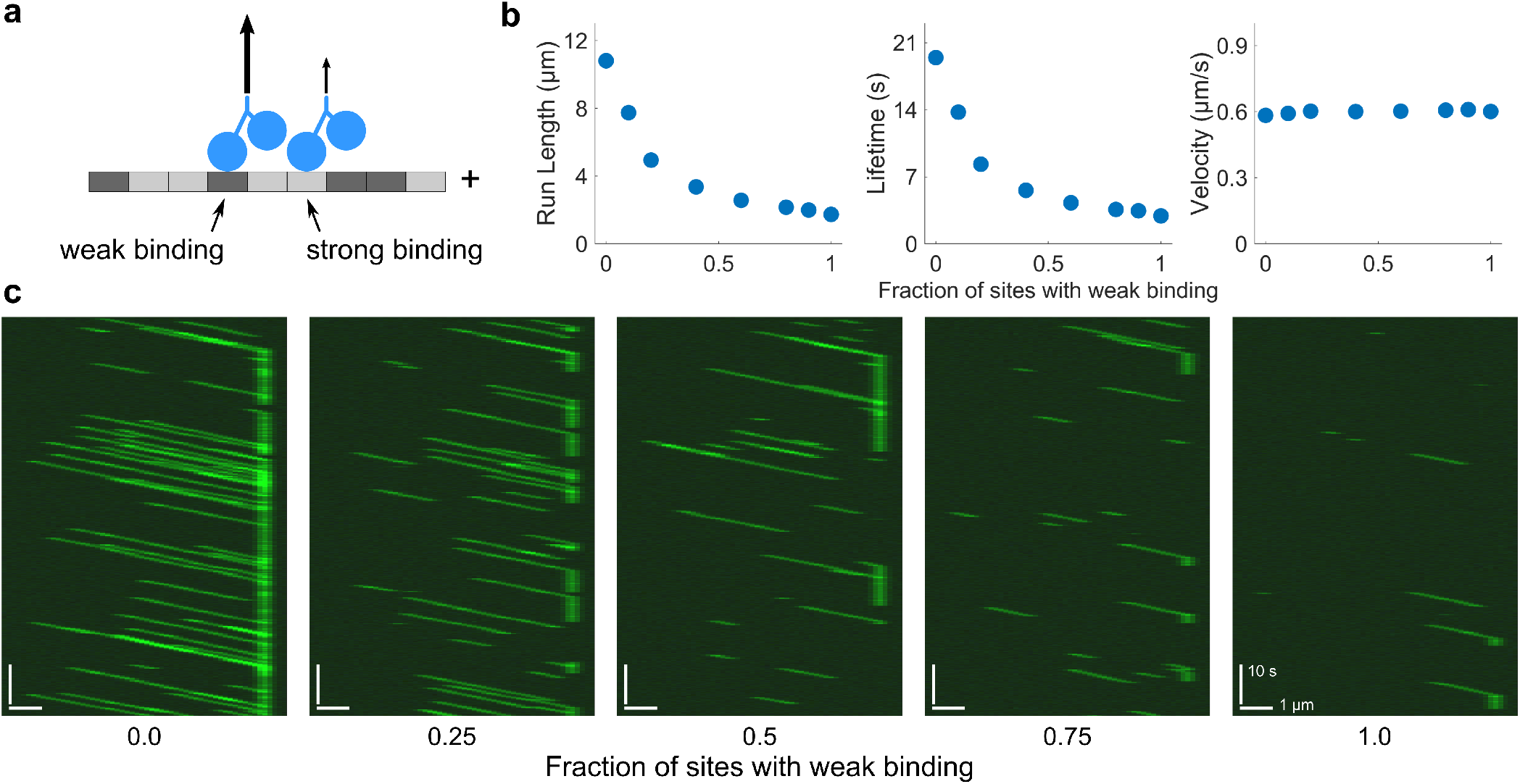
Effects of modeling heterogeneous tubulin with a mix of strong- and weak-binding sites. (a) Model schematic. A fraction of tubulin sites (dark grey) are weakly binding. (b) Plots of motor run length, lifetime, and velocity as a function of the fraction of sites with weak binding. Here, *ζ_i_* = 3 and *c* = 50 pM. Motor processivity has been increased by an order of magnitude from the reference parameter set. Data points are the average of five independent runs. Error bars correspond to the standard error of the mean and are typically smaller than the points. (c) Simulated kymographs with varying fraction of weak-binding tubulin with all motors fluorescently labeled. Here *c =* 150 pM.

### C. Heterodimeric motors

Some kinesin motors in cells are heterodimeric, for which the two motor heads that make up the dimer have different behavior [104–109]. Engineered kinesin heterodimers can been created with differences in catalysis between the two heads [113–115]. The explicit modeling of motor mechanochemistry in CyLaKS makes simulated heterodimers straightforward to model.

To illustrate this capability, we created a heterodimeric motor with one head that can diffuse when singly bound (Fig. 5). One (normal) head is immobile while singly bound, while the other (mutant) can diffuse along the lattice while singly bound. We assume that the normal catalytic head is always the first to bind and is leading. With our typical parameter set, the heterodimer spends about 5% of its time in a state that can diffuse. To see when one-head diffusion might have a significant effect, we decreased the effective concentration of the second motor head by a factor of 40. This increases the amount of time the motor spends in the a diffusible state to about 40% of the cycle. To keep overall motor processivity constant, we decreased the off rate of docked motors by a factor of 40.

**FIG. 5.**
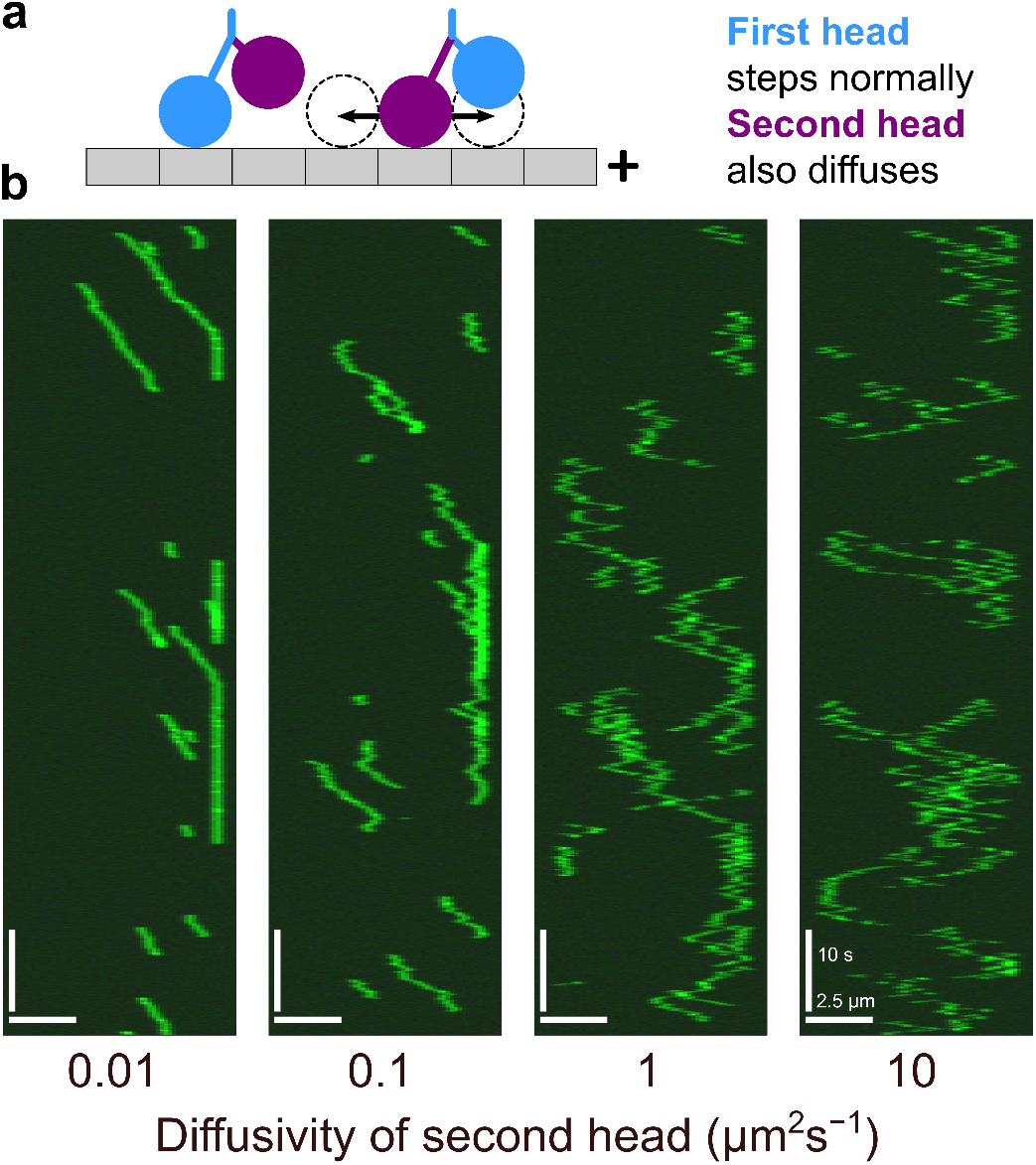
Kinesin heterodimer simulations. (a) Model schematic. Kinesin heterodimers are constructed of one normal head (blue) and one mutant head (purple) which diffuses while singly bound. (b) Simulated kymographs of kinesin heterodimer for *D* = 0.01, 0.1, 1, and 10 *μ*m^2^ s^−1^. All simulated molecules are fluorescently labeled.

With these parameters, we observe significant changes in motor activity as we increase diffusion coefficient of the second head (Fig. 5b). For low diffusion coefficient, processive runs still occur and motors pause at the ends of microtubules. For intermediate diffusion coefficient, the effects of diffusion are noticeable as fluctuations about the mean velocity during a processive run. For high diffusion coefficient, both processive movement and end pausing are reduced by the diffusion.

### D. Filament separation

In assemblies of many microtubules such as the mitotic spindle and reconstituted microtubule bundles, the spacing between microtubules can vary due to interactions defined by crosslinking motors and MAPs [99, 116–119]. While crosslinking motors and crosslinkers are typically 30-50 nm long [99, 120], the lateral separation between the surface of microtubules in the fission yeast spindle is typically 5-15 nm [116, 118]. This suggests that additional physical effects beyond just the length of motors or crosslinkers may be important for microtubule spacing in the spindle.

We developed a model of crosslinker-mediated control of the spacing of a pair of microtubules and studied it both analytically and in CyLaKS (Fig. 6). Two filaments can change their lateral separation *h* but are otherwise fixed in position. A fixed number of crosslinkers can diffuse along the lattice, and force exerted by the crosslinkers determines the filament separation. Crosslinkers that bind at an angle between the filaments are typically stretched (Fig. 6a), inducing force that pull the filaments closer together than the crosslinker length.

**FIG. 6.**
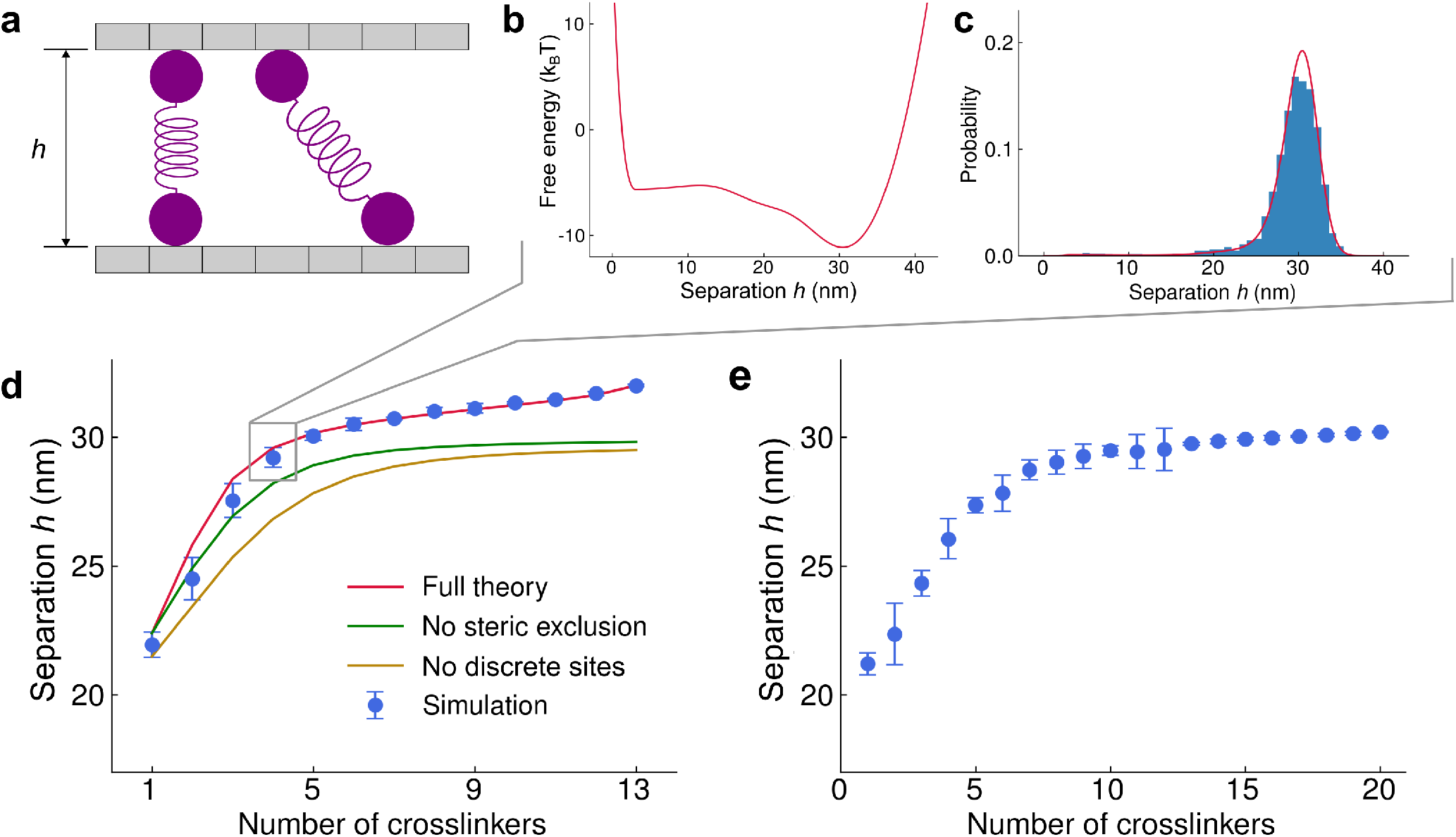
Change in microtubule separation with varying crosslinker number. (a) Schematic of the model. The microtubule separation *h* varies due to forces from crosslinkers. (b) Plot of free energy as a function of microtubule separation for 13-site microtubules with 4 32-nm-long crosslinkers, from semi-analytic theory. (c) Probability as a function of microtubule separation for 13-site microtubules with 4 32-nm-long crosslinkers from semi-analytic theory (red) and simulation (blue). (d) Average microtubule separation as a function of number of crosslinkers for 13-site microtubules, comparing full semi-analytic theory (red), simulation (blue), and theory neglecting steric exclusion (green) and both steric exclusion and discrete lattice sites (gold). (d) Average microtubule separation as a function of number of crosslinkers for 100-site microtubules.

For relatively short microtubules (up to 20 sites), the free energy of crosslinker binding and average separation can be computed by explicit enumeration of all possible binding states. To do this computationally, we first insert a single crosslinker in all possible binding states and note the crosslinker spring energy of each as well as possible binding states of additional crosslinkers. Then a second crosslinker is added and the procedure is iteratively continued to many crosslinkers, while ignoring crosslinker permutations. In this procedure we enforce steric interactions so that only one crosslinker can bind to each lattice site, and crosslinker crossing is not allowed. The crosslinker partition function is then

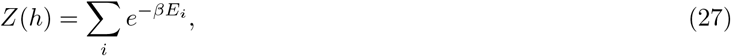

where *E_i_* is the crosslinker spring energy for a particular configuration *i* and the sum is over all possible arrangements of crosslinkers for a given separation *h*. The total free energy is then

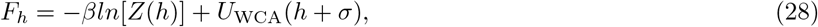

where *U*_WCA_ is the Weeks–Chandler–Anderson repulsive potential between microtubules, which acts on the microtubule centers separated by a distance *h + σ*, where *σ* is the microtubule diameter (Fig. 6b). The average separation is then

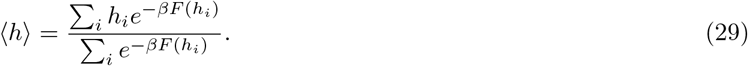

Simulations in CyLaKS accurately sample the Boltzmann distribution as microtubule separation varies (Fig. 6c). Here we used crosslinkers 32-nm long, as found for PRC1 [99]. When we vary crosslinker number for microtubules with 13 or 100 sites, we find that microtubule separation is typically smaller than 32 nm, with a particular drop for smaller crosslinker number (Fig. 6d,e). Altering the model to neglect crosslinker steric exclusion or discrete sites along the microtubule tends to predict slightly lower separation (Fig. 6d).

## IV. DISCUSSION

We developed the the Cytoskeleton Lattice-based Kinetic Simulator (CyLaKS) to facilitate modeling of cytoskeletal systems in which spatiotemporal changes and heterogeneity of the filament, motors, and/or associated proteins are significant. We built on previous theory and modeling of single motor mechanochemistry [80–86] by incorporating a model of motor ATP hydrolysis into a simulation with many interacting motors. This extends the approach of existing cytoskeletal simulation packages [66–69] to include a more detailed motor model. CyLaKS also implements detailed-balance in binding kinetics and movement, to model force-dependent protein binding/unbinding and diffusion. It is also designed to model spatiotemporally varying interactions and structure, such as filament lattice heterogeneity and short- and long-range interactions between motors. The framework is flexible and extensible, making it straightforward to elaborate the model.

In this paper we have shown examples of problems that CyLaKS can model. End-tags of kinesin-4 motors can form due to long-range interactions between motors. These end-tags depend on microtubule length, and dynamically adjust if the microtubule is ablated into two filaments. Implementing a heterogeneous microtubule lattice with a randomly located fraction of weak binding dimers shows that motor run length and lifetime decrease as the fraction of weak-binding sites increases, because motors unbind more quickly from the weak sites. Because CyLaKS is designed to model explicit motor stepping, it is straightforward to model an artificial motor heterodimer in which one head is fixed in place when singly bound, while the other head can diffuse. The overall motor movement then shows a crossover from directed runs to diffusion as the second head diffusion coefficient increases. Finally, we used both CyLaKS and analytic theory to show that crosslinker forces can lead to an equilibrium microtubule separation shorter than the croslinker length.

CyLaKS can be used to model motor and crosslinker behavior that is spatiotemporally altered. Problems including short-range interactions between motors [37, 38] and motor response to patchy obstacles [39] are straighforward to simulate. Closely related are changes in motor behavior due to crowding, both crowding along the filament lattice [40–45, 54] or due to crowders in solution [55], and motor direction switching [50–54]. Changes in the filament lattice, for example due to heterogeneous isoforms or post-translational modification [110–112] or lattice structure changes or defects [59–64] can be modeled in CyLaKS.

Extensions to CyLaKS to include additional mechanisms are straightforward and would be of interest in future work. The decentralized structure of CyLaKS means that just a single new file that defines a custom class is required, along with modifying the appropriate manager file to handle the class. Extensions to CyLaKS that would be of particular interest are incorporating microtubule dynamic instability, flexibility, and multiple protofilaments.

## Acknowledgements

We thank Matthew Glaser, Dick McIntosh, and Jeffrey M. Moore for useful discussions. We thank Radhika Subramanian and Sithara Wijeratne for useful discussions as well as access to unpublished data. This work was funded by NSF grant DMR-1725065 and NIH grant R01GM124371. Simulations used the Summit supercomputer, supported by NSF grants ACI-1532235 and ACI-1532236.

